# Dynamics of asymmetric membranes and interleaflet coupling as intermediates in membrane fusion

**DOI:** 10.1101/2022.07.31.502218

**Authors:** Marcos Arribas Perez, Paul A. Beales

## Abstract

Membrane fusion is a tool to increase the complexity of model membrane systems. Here, we use silica nanoparticles to fuse liquid-disordered DOPC giant GUVs and liquid-ordered DPPC:Cholesterol (7:3) GUVs. After fusion, GUVs display large membrane domains as confirmed by fluorescence confocal microscopy. Laurdan spectral imaging of the membrane phases in the fused GUVs shows differences compared to the initial vesicles indicating some lipid redistribution between phase domains as dictated by the tie lines of the phase diagram. Remarkably, using real-time confocal microscopy we were able to record the dynamics of formation of asymmetric membrane domains in hemifused GUVs and detected interleaflet coupling phenomena by which the DOPC-rich liquid-disordered domains in outer monolayer modulates the phase state of the DPPC:Cholesterol inner membrane leaflet which transitions from liquid-ordered to liquid-disordered phase.. We find that internal membrane stresses generated by membrane asymmetry enhance the efficiency of full fusion compared to our previous studies on symmetric vesicle fusion. Furthermore, under these conditions, the liquid disordered monolayer dictates the bilayer phase state of asymmetric membrane domains in >90% of observed cases. By comparison to the findings of previous literature, we suggest that which monolayer phase dominates the bilayer properties could be a mechanoresponsive signalling mechanism sensitive to the local membrane environment.

**Statement of Significance:** Natural biomembranes are highly asymmetric in lipid composition between the two leaflets of its bilayer structure. While the majority of membrane biophysics studies are conducted on symmetric lipid bilayers, it has become increasingly apparent that these asymmetric lipid compositions have a strong impact on the overall properties and behaviours of the membrane. However, achieving controlled membrane asymmetry in artificial model membranes is non-trivial, making the study of related biophysical mechanisms challenging. Here, we present an experimental framework to image the dynamics of asymmetric membrane domains formed as intermediates during membrane fusion of giant unilamellar vesicles (GUVs) triggered by silica nanoparticles. We directly observe interleaflet coupling of the phases in asymmetric membranes, where the liquid disordered domains most commonly dominate. By comparison with other literature on asymmetric membranes, this may indicate that the phase state of asymmetric membranes can be mechanoresponsive to its local environment. Furthermore, enhanced efficiency of full fusion of these GUV membranes indicates that stresses generated by membrane asymmetry can promote the formation of the full fusion pore during vesicle fusion processes.

## INTRODUCTION

Biological membranes are formed by a myriad of different lipids which provide them with particular biophysical properties and functionality (1). The organization of lipids within the membrane can lead to lateral heterogeneities where liquid ordered (*L*_*o*_) and liquid disordered (*L*_*d*_) phases separate into coexisting membrane domains (1-3). In cell membranes, lipid domains are transient and nanoscopic (<200 nm), hence difficult to characterize experimentally (3-6). For this reason, biomimetic model membranes, such as giant unilamellar vesicles (GUVs) and giant plasma membrane vesicles (GPMVs), which can display larger and more stable microdomains, have been extensively employed to study the lateral organization of lipids within phase separated membranes (1).

In addition to lateral heterogeneities, biological membranes also exhibit differences in the lipid content of each leaflet. This membrane asymmetry is conserved across organisms of all living kingdoms and plays a fundamental role in numerous cellular functions (7,8). Early studies proved the asymmetric distribution of just phospholipid headgroups (9,10), but Lorent *et al*. recently demonstrated that the plasma membrane of human erythrocytes is also asymmetric in the degree of saturation of the acyl chains, with the inner leaflets enriched in lipids with polyunsaturated acyl chains (11). Membrane asymmetry has represented a problem to understand the formation of *L*_*o*_ domains in the plasma membrane, since they exist in both leaflets but only the outer monolayer presents the appropriate lipid composition to form such structures (12,13). The reason behind the formation of bilayer domains is that the two monolayers are coupled, so one can modulate the physical state of the other and thus determine the physicochemical properties of the membrane bilayer (4,6,13). Nevertheless, the mechanisms governing the interleaflet coupling phenomena are still not clear and might involve many different physicochemical parameters such as line tension, surface tension at the bilayer midplane, acyl chain interdigitation, membrane fluctuations, curvature and differential lateral stress (4,6,7).

Commonly, studies on lipid lateral organization use GUVs prepared from solutions containing a mixture of the desired lipids to display phase separation. Similarly, most investigations of membrane asymmetry rely on model membranes directly produced with different lipid compositions in each leaflet. However, alternative methods to induce phase separation in initially homogeneous membranes can provide a more precise control over the final membrane composition and a better understanding on the dynamic processes behind the lipid organization within the membrane. Membrane fusion represents a potential strategy to fulfil this purpose. Bezlyepkina *et al* used electrofusion to generate three component GUVs from simpler vesicles and determine the tie lines of ternary lipid mixtures in the final GUV (14). Additionally, membrane phase separation can also be of great importance in synthetic biology as a spatiotemporal regulation system of membrane-protein interactions and membrane remodeling processes. For instance, Dreher *et al* showed that phase separation can lead to the division of a *L*_*d*_-*L*_*o*_ GUV into two single-phased daughter vesicles, one formed by the disordered phase and the other by the ordered phase (15). They then used DNA-triggered membrane fusion to induce phase separation in the daughter GUVs and repeat the division process (15). Other studies have also employed fusion-based strategies to generate asymmetric GUVs (16,17).

Previously, we showed that silica nanoparticles (SiO_2_ NPs) of 30 nm diameter effectively mediate fusion of DOPC GUVs by imposing a balance between membrane curvature and increased lateral tension in the membranes (18). Here, we use the same type of SiO_2_ NPs to mediate membrane fusion between *L*_*d*_ and *L*_*o*_ GUVs with the aim to generate phase separated GUVs. The fusions between *L*_*d*_ and *L*_*o*_ GUVs show mechanistic differences compared to fusions between two DOPC GUVs indicating that differences in the properties of each fusing membrane might help to promote the opening of the final fusion pore. Moreover, by recording the fusion process we visualize changes in lipid lateral organization, asymmetric domain formation and interleaflet coupling phenomena experienced by hemifused GUVs.

## MATERIALS AND METHODS

### Materials

Lipids 1,2-dioleoyl-sn-glycero-3-phosphocholine (DOPC), 1,2-dipalmitoyl-sn-glycero-3-phosphocholine (DPPC), cholesterol (Chol) from ovine wool, and 1,2-dioleoyl-sn-glycero-3-phosphoethanolamine-N-(lissamine rhodamine B sulfonyl) (ammonium salt) (Rh-DOPE) were purchased from Avanti Polar Lipids Inc. (Alabaster, USA). 3,3’-dioctadecyloxacarbocyanine perchlorate (DiO), and 6-Dodecanoyl-2-Dimethylaminonaphthalene (Laurdan) were purchased from ThermoFisher Scientific Ltd. (Loughborough, UK). Naphtho[2,3-a]pyrene (naphthopyrene) was obtained from Tokyo Chemical Industries UK Ltd (Oxford, UK). Silica nanoparticles LUDOX TM-50, HEPES, glucose, NaCl and indium titanium oxide (ITO) coated glass slides (surface resistivity 8–12 V sq-1) were purchased from Sigma-Aldrich Co. (Gillingham, UK). Microscope μ-slide 8 well glass bottom chambers (Ibidi GmbH) were purchased from Thistle Scientific Ltd (Glasgow, UK).

### Electroformation of GUVs

GUVs were prepared using the electroformation method (19). First, 15 µl of the desired lipid mixture in chloroform (0.7 mM) were deposited as a thin layer over the conductive side of two ITO coated glass slides and then dried under a nitrogen stream. The lipid mixtures used were DOPC and 0.5 mol% Rh-DOPE for the *L*_*d*_ GUVs; and DPPC:Chol (7:3 mol ratio) with 1 mol% of naphthopyrene for the *L*_*o*_ GUVs. The ITO slides were then assembled into an electroformation chamber each in contact with a copper tape and separated by a 1.6 mm Teflon gasket. The chamber was filled with a 300 mM sucrose solution (300 mOsm/kg) and connected to a function generator to apply an AC electric field. The frequency of the electric field was set at 10 Hz and the voltage was gradually increased from 1 V peak-to-peak (Vpp) to 5 Vpp over 15 minutes and maintained at 5 Vpp for two hours. Finally, the frequency was gradually reduced to 0.1 Hz over 10 minutes to facilitate the closure and detachment of GUVs from the slide. DOPC GUVs were prepared at room temperature whereas DPPC:Chol (7:3) GUVs were prepared at 50 °C (above melting temperature of DPPC (*T*_*m*_ of DPPC = 41 °C). After electroformation, GUVs were suspended in isotonic buffer containing 20 mM HEPES and 150 mM NaCl (300 mOsm/kg) adjusted to pH 7.4. The osmolality of the inner sucrose solution and the external buffer was measured with a 3320 Micro-Osmometer (Advanced Instruments, Norwood, UK).

### Confocal imaging and analysis

Imaging was performed with a Zeiss LSM-880 inverted laser scanning confocal microscope with a Plan-Apochromat 40x/1.4 Oil DIC M27 objective lens (NA = 1.4). To detect the membrane phase DOPC GUVs were labelled with Rh-DOPE which partitions into liquid disorder phase and DPPC:Chol (7:3) GUVs with naphthopyrene (Npy) which partitions into liquid ordered phase. 100 µl of each population of GUVs were deposited into a well of a microscope slide previously passivated with a solution of 5% BSA for 10 minutes followed by rising with Mili-Q water. Once the GUVs were sunk in the bottom of the well, 25 µg/ml SiO_2_ NPs were carefully added to the sample and we acquired time series to record fusion events in real-time. Npy and Rh were excited with a 405 nm diode laser and a 561 nm diode pumped solid state (DPSS) laser, respectively. The emission of Npy was recorded between 410 nm and 500 nm and the emission of Rh between 566 nm and 630 nm. Image analysis was performed with Fiji (ImageJ) and plotting and statistical analysis were done using Origin Pro. Real-time Tile scanning was used to image large areas of the sample and the proportion of phase separated GUVs in the sample after exposure to 25 µg/ml SiO_2_ NPs was counted manually.

### Laurdan Spectral imaging

GUVs labelled with 0.5 mol% Laurdan were prepared by electroformation. The spectral images are acquired using the lambda mode of the Zeiss LSM880 confocal laser microscope, which allows split the fluorescence emission spectrum of a dye into up to 34 spectral intervals recorded in individual detection channels. Laurdan was excited with the 405 nm laser and the fluorescence detection range was set between 410 nm and 550 nm with a spectral step of 8.9 nm per channel. Snapshots of Laurdan labelled GUVs were acquired before and after exposure to NPs. The images were analyzed with a Fiji plugin developed by Sezgin *et al*.*(20)*, setting 440 nm and 490 nm as maximum emission wavelengths to calculate the GP values using the following equation:

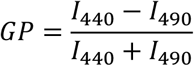

where *I*_*440*_ and *I*_*490*_ are the fluorescence intensity at 440 nm and 490 nm, respectively.

## RESULTS AND DISCUSSION

### SiO_2_ NPs mediate fusion between *L*_*d*_ and *L*_*o*_ GUVs

We first tested the ability of 30 nm SiO_2_ NPs to promote fusion between a *L*_*d*_ GUV and a *L*_*o*_ GUV. Two populations of GUVs were prepared using electroformation, one made of DOPC (*L*_*d*_ GUVs) and the other composed of DPPC and cholesterol (Chol) in a 7:3 mol ratio (*L*_*o*_ GUVs). The two populations of GUVs were mixed in a 1:1 volume ratio and incubated with 25 µg/ml of SiO_2_ NPs. Experiments were performed at room temperature, below the melting temperature of DPPC (*T*_*m*_= 41°C). To visualize the reorganization of the lipids in the membrane during the fusion process, we incorporated Rh-DOPE (0.5 mol%) to the *L*_*d*_ GUVs and naphthopyrene (Npy) (1 mol%) into the *L*_*o*_ GUVs. These dyes are used as membrane phase markers as Rh-DOPE preferentially partitions into the *L*_*d*_ phase of the membrane and Npy into *L*_*o*_ domains (21).

Using real-time confocal microscopy, we observe that 30 nm SiO_2_ NPs promote fusion between a *L*_*d*_ GUV and a *L*_*o*_ GUV. Importantly, once fused, the membrane of the new GUV lowers its free energy by remaining phase separated, displaying large *L*_*d*_ and *L*_*o*_ microdomains (Fig. 1, Supplementary movie 1). Our observations are consistent with previous studies on membrane phase separation in DOPC:DPPC:Chol GUVs (22). Considering that the initial GUVs are similar in size, the final membrane composition of the fused GUV is predicted to be close to 50:35:15 DOPC:DPPC:Chol. At room temperature, this ternary lipid composition is in the liquid-liquid immiscibility region of the phase diagram, where *L*_*d*_ and *L*_*o*_ domains coexist in the membrane (22).

**Figure 1.**
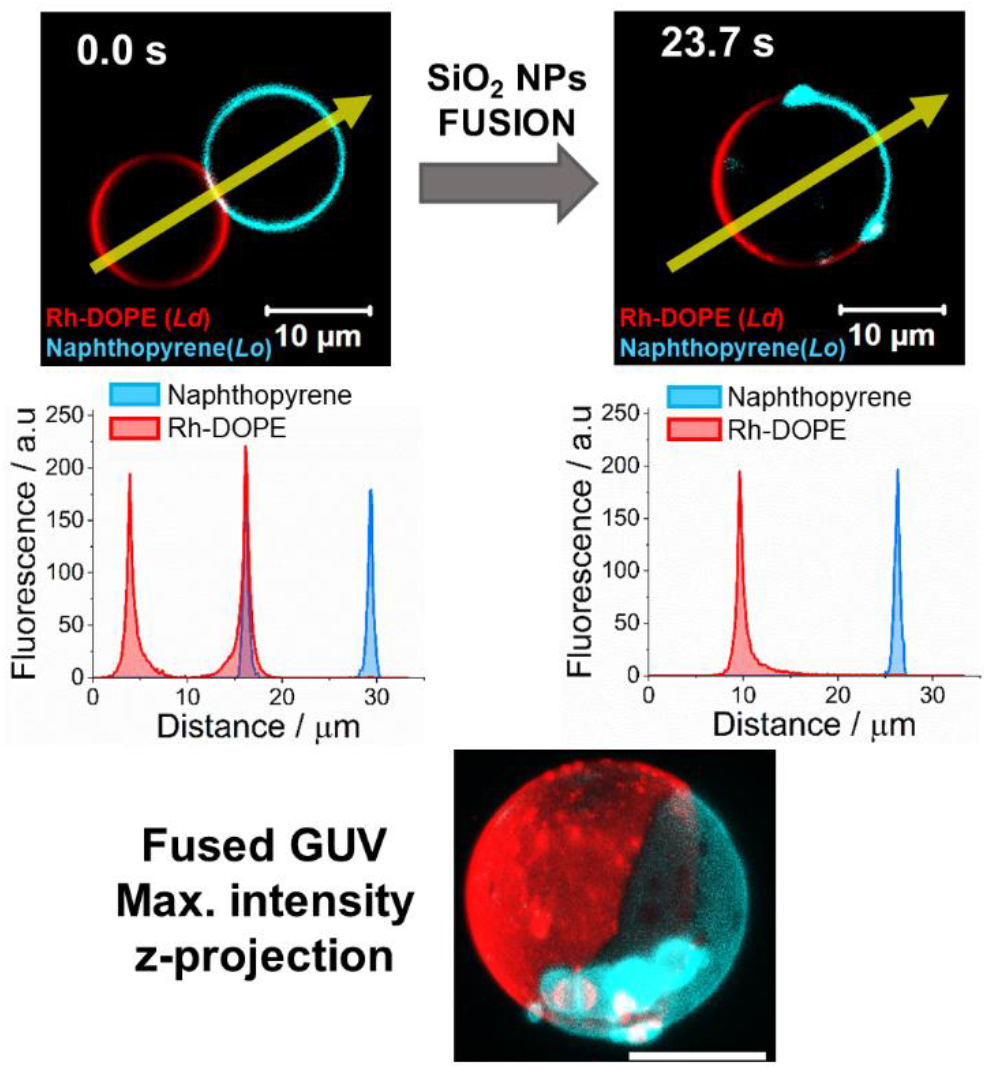
Confocal microscopy images of *L*_*o*_ and *L*_*d*_ GUVs before and after fusion. *L*_*o*_ DPPC/Chol (7:3) GUV is labelled with naphthopyrene (Npy) (cyan) and *L*_*d*_ DOPC GUV is labelled with Rh-DOPE (red). After fusion Npy partitions into the *L*_*o*_ phase and Rh-DOPE into the *L*_*d*_ phase. Plots show the fluorescence intensity profile of Rh-DOPE (red) and Npy (blue) along the yellow arrow shown in the micrographs. Bottom image shows a 3-D maximum intensity projection of a phase separated fused GUV.

Further analysis of the membrane phase was performed using Laurdan spectral imaging. In this case, *L*_*d*_ GUVs and *L*_*o*_ GUVs were both labelled with 0.5 mol% Laurdan. In addition, 0.1 mol% Rh-DOPE was also added to the *L*_*d*_ GUVs. Control *L*_*d*_ GUVs show mean Laurdan GP values of −0.45 ± 0.02, while the GP values of *L*_*o*_ GUVs are much higher, peaking at 0.29 ± 0.03 (Fig. 2a), as expected for more ordered membranes (20). After fusion, the Laurdan GP map clearly shows two easily distinguishable regions in the membrane, which correspond to the *L*_*o*_ (positive GP values) and *L*_*d*_ (negative GP values) domains (Fig. 2b). Rh-DOPE fluorescence in the fused GUV matches perfectly with the region of the membrane displaying negative GP values, confirming their *L*_*d*_ nature (Fig. 2b). Compared to the control DOPC GUVs, the *L*_*d*_ domains of the fused GUVs show significantly higher (less negative) average GP values (GP= −0.10 ± 0.05), which indicates a denser lipid packing while the GP values observed in the *L*_*o*_ domains of the fused GUVs are slightly less positive (GP= 0.24 ± 0.04) than in the control DPPC:Chol GUVs (Fig. 2c). After fusion, the lipids laterally redistribute between membrane domains until an equilibrium lipid composition, determined by the tie lines of the phase diagram, is reached. Consequently, the *L*_*d*_ phase of the fused GUV is no longer a pure DOPC membrane but a DOPC-rich domain, and the same occurs with the *L*_*o*_ phase, which is enriched in DPPC and cholesterol but not exclusively formed by these lipids.

From tile scan images, we quantified the average proportion of phase separated GUVs generated after incubating the vesicles with the SiO_2_ NPs to be 11.25 ± 3.89% (Fig. S1 a). This value does not show statistically significant difference with respect to the proportion of fused GUVs observed in samples of just DOPC GUVs in isosmotic conditions (18) (Fig. S1 a). These results demonstrate that the fusogenic activity of 30 nm SiO_2_ NPs is not limited to DOPC membranes but they maintain their ability to promote fusion when one of the membranes is stiffer, as in the case of DPPC:Chol GUVs.

In a recent study, we observed that the fusion mediated by these SiO_2_ NPs can follow three different pathways that we classified as direct full fusion, hemifusion-fusion and gentle membrane merging depending on the intermediate states that the GUVs adopt during the process (18). In the direct full fusion, the fusion occurred before any lipid transfer between the GUVs was detected, while the other two mechanisms involved lipid mixing indicative of hemifusion intermediate states. In the hemifusion-fusion processes, the hemifusion is followed by the opening of a fusion pore whereas in the gentle membrane merging events the fusion pore never opens and one of the GUVs is gradually absorbed by the other. From our results we did not see any preferential pathway with 35% of the fusions following the direct full fusion pathway, 30 % the hemifusion-fusion pathway and the remaining 35% undergoing gentle membrane merging (Figure S1 b). We proposed that the fusion pathway followed by the GUVs would be primarily determined by the membrane tension, so fusing GUVs that do not have enough tension to overcome the energy cost associated to the opening and expansion of a fusion pore would follow the gentle membrane merging pathway (18).

Here we recorded 32 confocal microscopy time-series of complete fusion events. The analysis of the movies showed that in just over half of the movies (53%, n=17) the GUVs experienced a direct full fusion (Fig. 1, Supplementary movie 1). In the other 15 observations (47%), we detected a hemifusion intermediate state before a fusion pore opens and the GUVs fuse. A more thorough analysis and discussion of the hemifusion processes will be provided later. Physically, the differentiation between the direct full fusion and the hemifusion-fusion pathways is one of kinetics. In the direct full fusion pathway, the full fusion pore opens on a rapid timescale where the hemifusion intermediate is not detectable with the temporal resolution of the time-resolved confocal imaging (frame rate = 3 Hz).

Interestingly, the gentle membrane merging pathway, which represented a 35% of the total fusions between two DOPC GUVs recorded in our previous study (18) (Figure S1 b), is completely suppressed when one of the GUVs has a *L*_*o*_ membrane. The reason for the absence of gentle merging membrane events is unclear. One possibility is that the transbilayer asymmetry at the hemifusion diaphragm, formed by a DOPC monolayer and a DPPC:Chol monolayer, might make the membrane more unstable or increase the membrane tension and thus lower the energy barrier to the eventual opening of the fusion pore. Using flicker spectroscopy, Elani *et al*. demonstrated that asymmetric GUVs, with one leaflet composed of DOPC and the other with POPC, show a significantly increased membrane bending rigidity compared to symmetric bilayers (23). Theoretical modelling and coarse-grained simulations have also suggested that imbalanced lipid packing in each leaflet of asymmetric membranes can generate a differential lateral stress which can have strong impact in the mechanical properties of the membrane, including membrane tension, bending rigidity and spontaneous curvature (24). During the fusion of *L*_*o*_ and *L*_*d*_ GUVs, the increase of membrane tension at the asymmetric hemifusion diaphragm would then provide the energy needed for the opening and expansion of a fusion pore. Our results therefore suggest an important role of the different lipid composition and phase state of each membrane in the opening of the full fusion pore in the last stage of the fusion process.

Notably, while the efficiency of opening of a full fusion pore is enhanced by asymmetric hemifused membrane intermediates, the kinetics of full fusion pore opening is not obviously changed. The relative proportion of fusion events that are categorised as direct full fusion and as hemifusion-fusion pathways remain comparatively equal, as also observed for the case of symmetric DOPC-DOPC GUV fusion (Figure S1 b).

### Hemifusion between *L*_*d*_ and *L*_*o*_ GUVs leads to membrane asymmetry and interleaflet coupling

In nearly half of the fusion events recorded (n=15), the opening of the fusion pore was preceded by a hemifusion intermediate state. This initially gives rise to a metastable asymmetric membrane diaphragm during the fusion process. Figure 3a (and figure S2a) shows a confocal microscopy time-lapse of two GUVs undergoing hemifusion along with fluorescence intensity data in different regions of their membranes. Maximum fluorescence intensity of rhodamine (*Rh*_*max*_) corresponds to regions of the membrane where both leaflets are formed mainly by *L*_*d*_ DOPC lipids, whereas in *L*_*o*_ bilayers show maximum fluorescence intensity of Npy (*Npy*_*max*_). If the dyes were present in only one of the membrane monolayers, their fluorescence intensity would be ∼50% lower than those maximum values.

**Figure 2.**
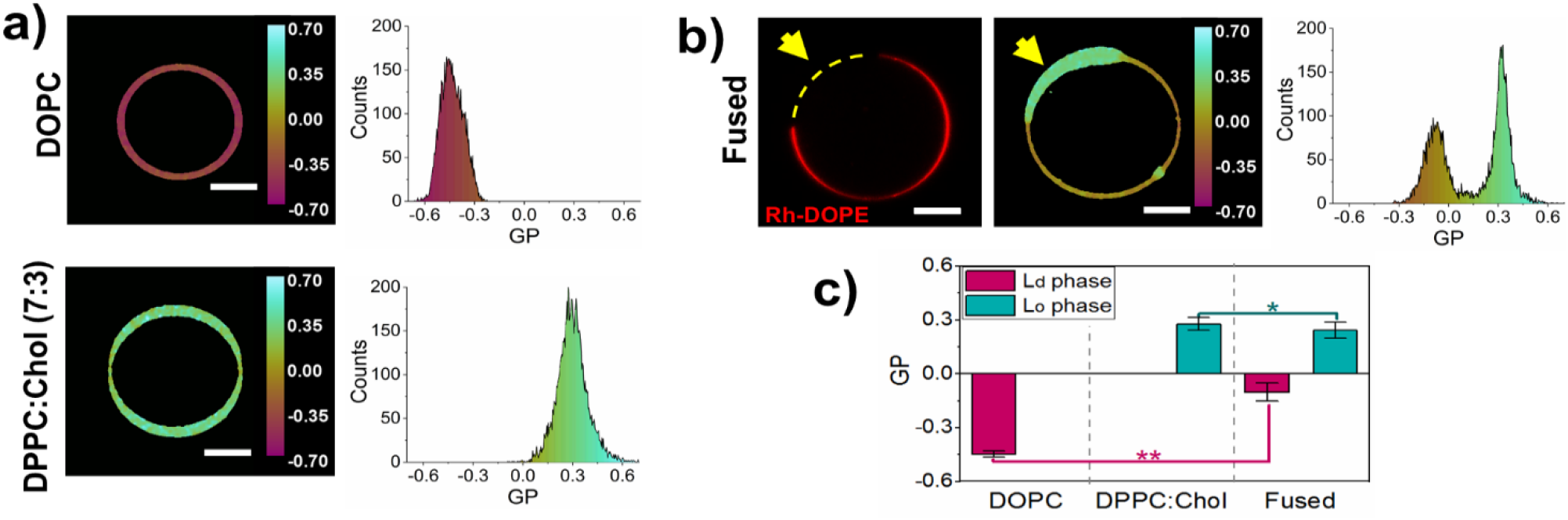
Laurdan GP of GUVs before and after fusion. a) Example of Laurdan GP maps and histograms of DOPC GUV (top) DPPC:Chol (7:3) GUV (down) GUVs. b) Example of Rh-DOPE fluorescence (left), Laurdan GP map (middle) and GP histogram (right) of fused GUV displaying phase separation. Rh-DOPE fluorescence matches with lower values of Laurdan GP (*L*_*d*_ phase) while it is excluded from the *L*_*o*_ domain which shows positive GP values. c) Bar plot of average GP values ± SD of *L*_*d*_ and *L*_*o*_ phase in DOPC GUVs (n= 26), DPPC:Chol (7:3) GUVs (n=20) and fused phase separated GUVs (n=26). The average GP value of the *L*_*d*_ phase is significantly less negative in the fused GUVs (GP= −0.10±0.05) than in the pure DOPC GUVs (GP= −0.45±0.02). The average GP of the *L*_*o*_ phase is slightly higher in the *L*_*o*_ GUVs (GP= 0.29±0.03) than in the fused phase separated GUVs (GP= 0.24±0.04). The statistical significance was tested using a one-way ANOVA with Bonferroni multiple comparisons test (*p ≤ 0.05, **p ≤ 0.01).

**Figure 2.**
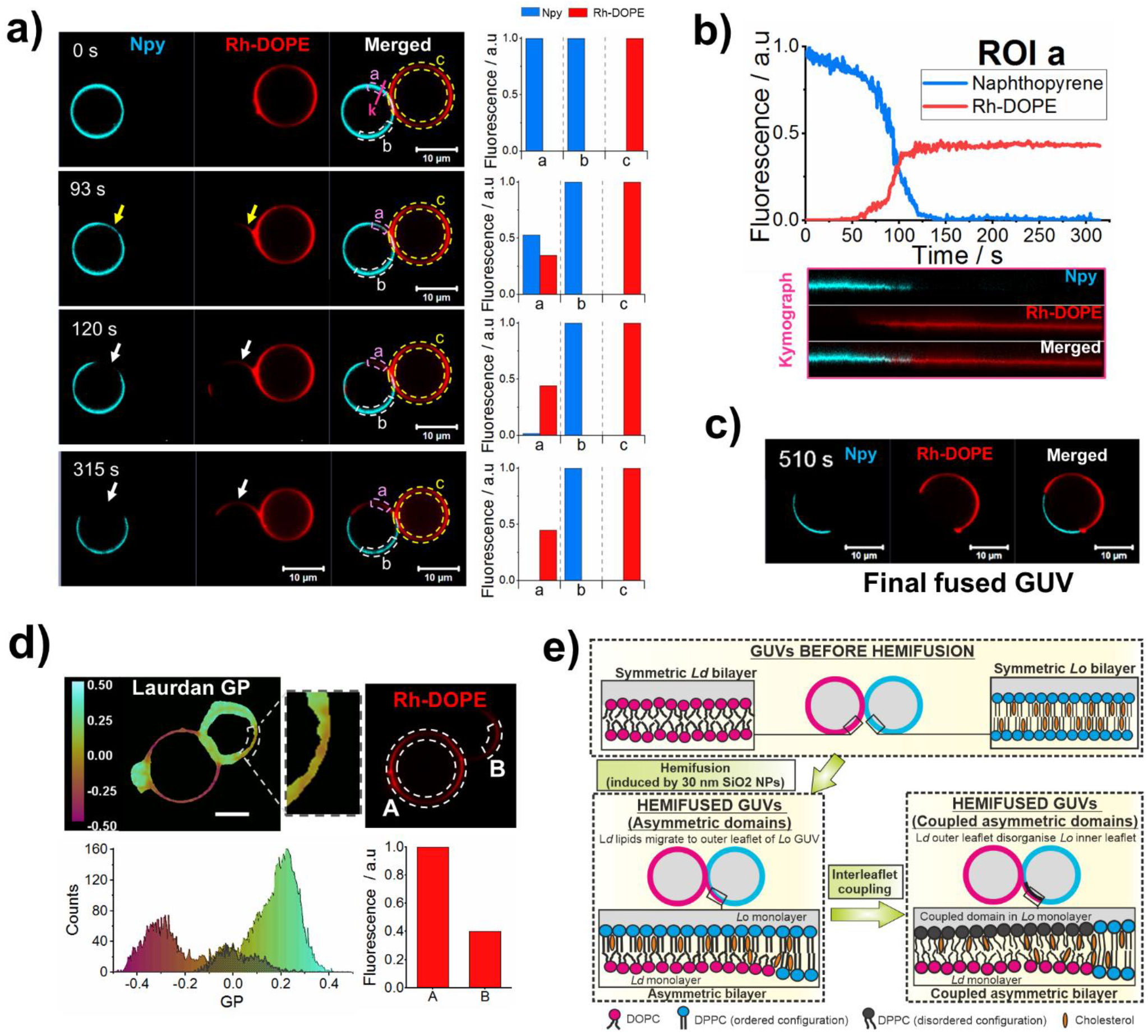
Asymmetric membrane domains and interleaflet coupling in hemifused GUVs. a) Confocal microscopy time-lapse images of GUVs during hemifusion. Naphthopyrene (Npy) (cyan) and Rh-DOPE (red) are used as *L*_*o*_ and *L*_*d*_ markers, respectively. Yellow arrows indicate regions of the membrane where both dyes are colocalised (asymmetric domains) and white arrows indicate regions of the membrane where the Npy has been excluded from the membrane. Bar plots show the normalised fluorescence intensity measured in the ROIs a, b and c shown in the merged channel. Fluorescence intensity in ROI b and ROI c corresponds to the maximum intensity of Npy and Rh-DOPE, respectively. ROI a shows an initial rise in Rh-DOPE intensity simultaneous to a reduction of Npy fluorescence to about half compared to its maximum (93 s micrograph). Later micrographs show a complete exclusion of Npy from that region of the membrane while the intensity of Rh-DOPE in ROI a remains close to 0.5. b) Normalised fluorescence intensity of Rh-DOPE (red line) and Npy (blue line) in ROI a against time. Kymographs show the fluorescence intensity over time at the pink line indicated as “k” in the 0 s micrographs in a). c) Phase separated GUV after full fusion. d) Laurdan GP map (left) and localisation of Rh-DOPE in the membrane (right) of a hemifused GUV. Zoomed region indicates a disordered domain within the initial *L*_*o*_ GUV that shows GP values (shaded in GP histogram) intermediate between the DOPC-rich membrane and the membrane enriched in DPPC and cholesterol. Fluorescence intensity of Rh-DOPE in ROI B denotes that the *L*_*d*_ marker is only present in one leaflet. e) Schematic representation of asymmetric domain formation and interleaflet coupling during hemifusion between a *L*_*d*_ GUV and a *L*_*o*_ GUV.

When the GUVs become hemifused, Rh-DOPE fluorescence emerges in a region of the *L*_*o*_ membrane. The Npy fluorescence intensity in the same membrane region is simultaneously reduced to about half compared to its fluorescence in the rest of the membrane, *Npy*_*max*_ (Fig. 3a, Fig. S2a, Supplementary movie 2). Assuming that only the outer leaflets of the GUVs are fused, the fluorescence intensities suggest that the *L*_*o*_ marker is only present in the inner membrane leaflet while the outer monolayer is enriched with DOPC lipids transferred from the opposite GUV, which induce a phase transition from *L*_*o*_ to *L*_*d*_ indicated by the rise of Rh-DOPE fluorescence. Hence, the structure in that region of the membrane would correspond to an asymmetric domain with an ordered DPPC:Chol inner leaflet and a disordered DOPC-rich outer monolayer (Fig. 3e). In all the events observed, it is the Rh-DOPE fluorescence that is transferred to a region on the *L*_*o*_ vesicle and never the other way around.

**Figure 3.**
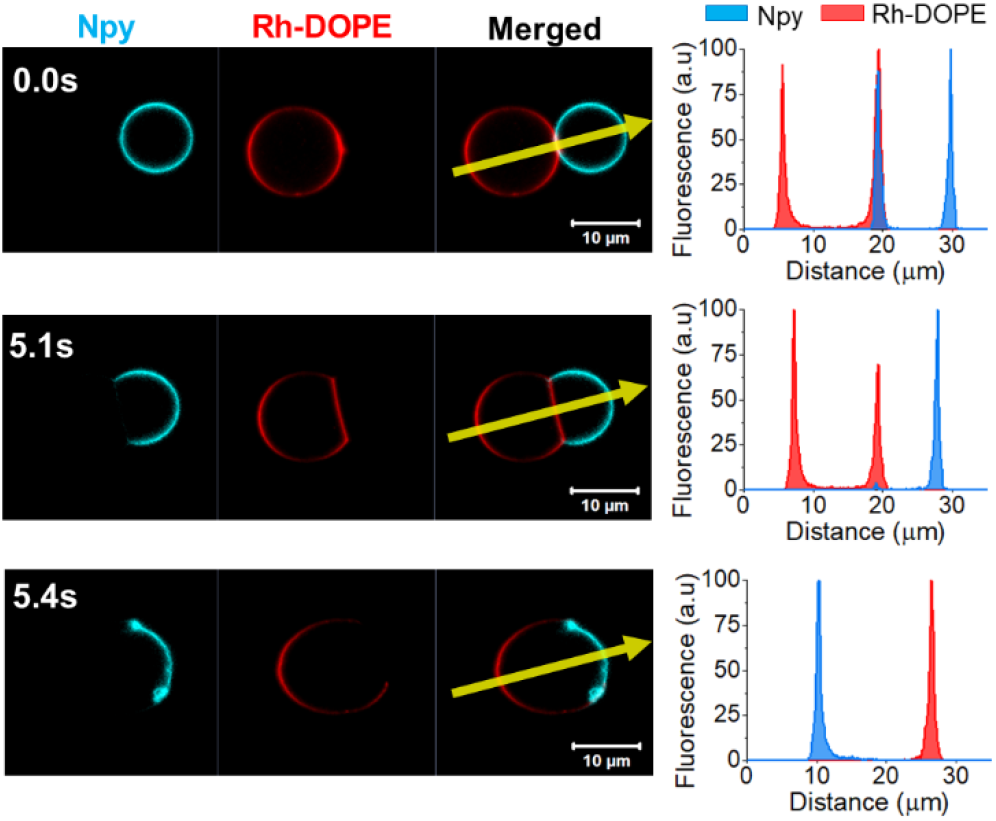
Confocal microscopy time-lapse micrographs showing naphthopyrene (Npy) exclusion from GUV-GUV interface prior to full fusion. Naphthopyrene (Npy) (cyan) and Rh-DOPE (red) are used as *L*_*o*_ and *L*_*d*_ markers, respectively. Plots show the fluorescence intensity profile of Npy (blue) and Rh-DOPE (red) across the yellow arrow indicated in the micrographs.

Strikingly, shortly after the formation of the asymmetric domain, we observe a complete exclusion of the *L*_*o*_ marker from the domain while the fluorescence intensity of Rh-DOPE remains close to 50% of *Rh*_*max*_ (Fig. 3a, Fig. S2a). The analysis of the fluorescence intensity dynamics in that membrane region clearly shows the transient colocalisation of the two dyes followed by a rapid the loss of Npy fluorescence (Fig. 3b, Fig. S2b). The exclusion of Npy from the asymmetric domain likely corresponds to an interleaflet coupling phenomenon by which a disorganisation in the DPPC:Chol inner monolayer is induced by the *L*_*d*_ outer leaflet, without noticeable interleaflet transfer of *L*_*d*_-prefering lipids, to create symmetry in the phase structure of the bilayer despite the compositional asymmetry being retained. The resulting structure would be an asymmetric *L*_*d*_ domain where Rh-DOPE molecules partition into the DOPC-rich outer leaflet while the inner leaflet is composed of DPPC and cholesterol in a disorganized configuration, which compels the Npy molecules to diffuse laterally and partition to regions where the lipids are more densely packed in an *L*_*o*_ phase (Fig. 3e). The loosening of the lipid packing within the asymmetric membrane domains was confirmed by Laurdan imaging (Fig 3d). The membrane asymmetry is only maintained while the GUVs are hemifused. Once the GUVs fully fuse, the membrane symmetry is restored and the location of the fluorescent dyes in one monolayer matches its location in the opposite monolayer (Fig.3c, Fig. S2c).

Two of the videos that we categorized as hemifusion show dynamical changes in the membrane phase at an apparently extended hemifusion diaphragm. In those events, we observe the complete exclusion of the *L*_*o*_ marker from the membrane region where the GUVs come into contact (Fig. 4, Fig. S3, Supplementary movie 3). The exclusion of Npy occurs simultaneously to an expansion and flattening of the contact region and a decrease of the relative fluorescence intensity of Rh-DOPE. The relative fluorescence intensity of the latter dye is however slightly higher (∼70%) than 50%. This fluorescence intensity data makes it difficult to confidently define the configuration adopted by the membrane. A possible scenario is that the observed structure is an expanded hemifusion diaphragm with a DOPC monolayer labelled with Rh-DOPE and a DPPC:Chol monolayer from which the Npy dye is excluded due to the interleaflet coupling. We speculate that the higher levels of Rh-DOPE fluorescence might be due to some extent of interleaflet lipid transfer, likely by a period of lipid flip-flop enhanced by membrane stresses during the initial hemifusion. The expanded hemifusion diaphragm dominated by the *L*_*d*_ phase shows a very short lifetime as it is formed just before the GUVs fully fuse. In these events, we do not detect migration of dyes from one GUV to the opposite. The dye exclusion and phase dynamics associated with it are restricted to the presumed hemifusion diaphragm and the *L*_*d*_ - *L*_*o*_ phase boundaries locate at the edges of it, where the bilayer at the hemifusion diaphragm meets the bilayers of the *L*_*d*_ and *L*_*o*_ GUVs, forming a Y-like junction.

**Figure 4.**
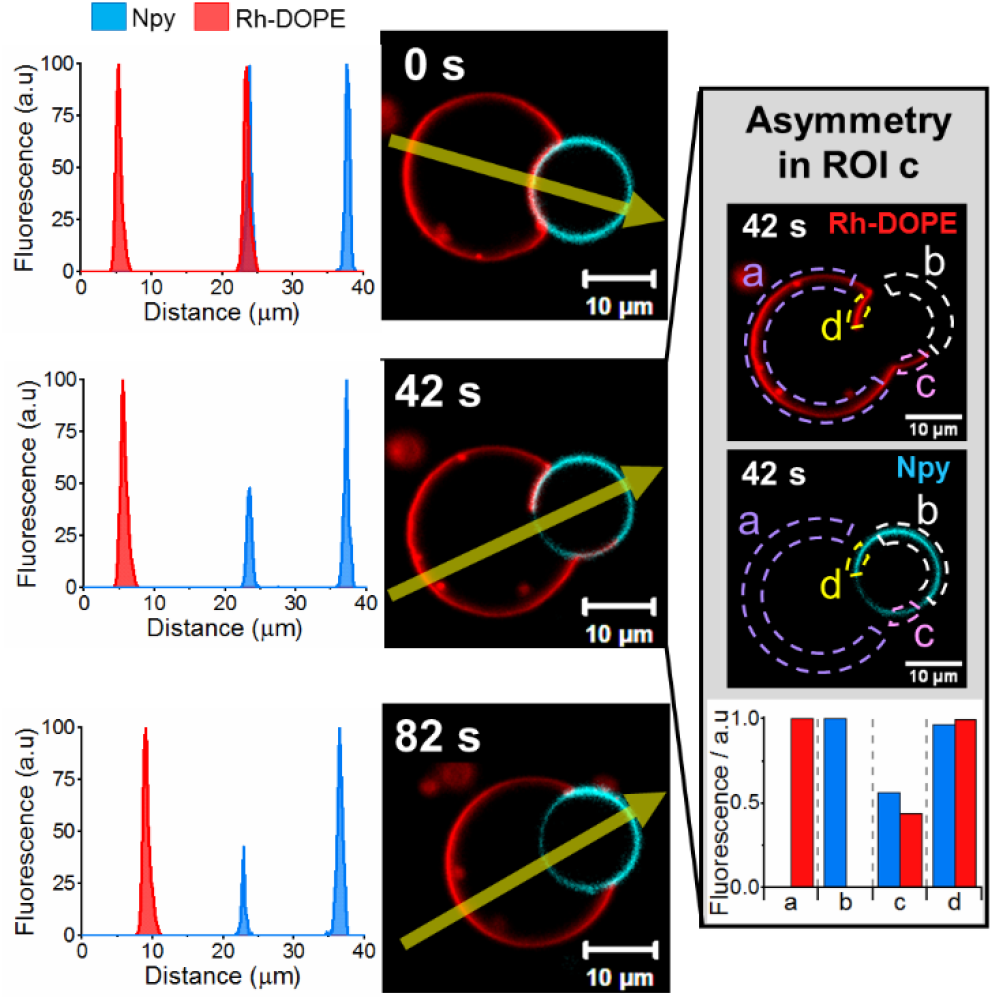
Confocal microscopy micrographs showing Rh-DOPE exclusion from hemifusion diaphragm. The dye exclusion advances gradually from one edge to the other. The plots in the left show the fluorescence intensity profile of Npy and Rh-DOPE across the yellow arrow indicated in the micrographs. Inset shows micrographs of the separated imaging channels at 42 s (Rh-DOPE on the top and Npy in the middle) and a detailed analysis of the fluorescence intensity (bottom bar plot) of each imaging channel in different regions of the membrane as shown in the micrographs. The fluorescence intensity in ROI c shows a membrane domain where the relative intensity of each dye is approximately 50% of their maximum fluorescence intensity (ROI b=Npymax; ROI c=Rhmax) that likely indicates membrane asymmetry.

In our experiments we only detected one event where the *L*_*d*_ marker is the one excluded from the membrane at the GUV-GUV interface, i.e. only 6% of the observed hemifusion events (Fig. 5, Supplementary movie 4). The exclusion of the *L*_*d*_ marker starts at one edge of the region where the two GUVs are in contact and it gradually advances towards the other edge. In this event, the hemifusion diaphragm does not become flattened but the *L*_*o*_ GUV pushes and bends the *L*_*d*_ GUV. In addition, the fusion process is not completed but the GUVs remain adhered by the hemifusion diaphragm (Note that as the fusion is incomplete, this movie was not counted within the 32fusion events mentioned earlier in the text). The deformation of the *L*_*d*_ GUV by the *L*_*o*_ GUV might indicate that the former vesicle is not tense enough for the fusion pore to open and expand. We also observe a lipid transfer from the *L*_*d*_ to the *L*_*o*_ GUV, which leads to the formation of an apparent asymmetric domain in the *L*_*o*_ GUV, similar to what we observed in the other hemifusion events, where the Npy relative fluorescence drops down to nearly 50% and the Rh fluorescence rises reaching about 50% of its value in the *L*_*d*_ GUV (Fig.5). The asymmetric domain is transient and we do not see interleaflet coupling but the Rh-DOPE-labelled membrane retreats to its original GUV and the membrane recovers its *L*_*o*_ symmetric appearance. Eventually, the phase boundaries locate at the edges of the hemifusion diaphragm and the membranes seem to reach an equilibrium state.

To our knowledge, this is the first time that asymmetric domain formation and interleaflet coupling are imaged in real-time in GUVs and during fusion processes. Enoki and Feigenson developed a method by which they created asymmetric GUVs *via* hemifusion of the vesicles with a supported lipid bilayer (SLB) (16). Using this method, they saw that after hemifusion, the phase separated inner monolayer of the GUVs induces phase separation in an otherwise *L*_*d*_ outer monolayer, however they only showed the final membrane configuration but the process leading to the interleaflet coupling is not revealed (16,17). In their system, the *L*_*o*_ domains modulate the lipid order in the opposing leaflet whereas our results show the opposite tendency. Nevertheless, they also see that the *L*_*o*_ domains in the asymmetric membrane are less ordered than the symmetric *L*_*o*_ phase. Previous studies on asymmetric lipid membranes show disparity regarding which phase dominates the final bilayer phase. For instance, *L*_*o*_ domains in the outer leaflet of SLBs have been seen to induce phase separation in the inner leaflet (25,26). However, the substrate where SLBs are formed can have a significant influence on the dynamical behavior of the lipids in the inner leaflet. Chiantia and London showed that long-chain sphingomyelin (SM) in the outer leaflet of vesicles decreases the lateral diffusion of DOPC lipids in the inner monolayer (27). They also saw that the interleaflet coupling is stronger when the PC lipids in the inner monolayer have one saturated acyl chain and attributed the coupling effect to the interaction between acyl chains at the bilayer midplane (27).

On the contrary, a disorganization effect of ordered domains by disordered lipids in the opposite leaflet, analogous to what we observe in most of our results, was reported by Heberle in asymmetric LUVs where the outer leaflet was composed of DPPC and the inner leaflet of POPC (28). Using small-angle neutron scattering (SANS), they observed that the *L*_*d*_ inner monolayer induced a significant decrease in the lipid packing density of the gel domains in the outer leaflet (28). Similarly, St. Clair *et al* have shown that while symmetric LUVs formed by a mixture of SM, POPC and cholesterol are phase separated, in asymmetric LUVs with SM:POPC:Chol in the outer leaflet but a POPC:Chol inner monolayer the formation of *L*_*o*_ domains is completely suppressed (29). Wang and London also reported the ability of DOPC:Chol *L*_*d*_ inner leaflet of asymmetric LUVs to destabilize and destroy ordered domains in the outer monolayer containing SM (30). This disorganization of the outer leaflet *L*_*o*_ domains is however less pronounced when its content in SM increases (30). Collins and Keller studied asymmetric pore spanning planar membranes formed by a *L*_*d*_ inner leaflet and an outer leaflet with a lipid composition within the *L*_*d*_*-L*_*o*_ coexistence region of the phase diagram and showed that increasing or decreasing the fraction of high-*T*_*m*_ lipid in the outer monolayer induces or suppresses, respectively, the formation of bilayer domains (31).

Our observations (and the studies mentioned above) confirm that, in bilayers of asymmetric composition, the phase of one leaflet can alter the phase state of the opposite leaflet to create phase symmetry in a bilayer domain. Nevertheless, the mechanisms that dictate which leaflet and which phase dominates over the other still remain elusive. The type of model membrane employed as well as the varying lipid composition and experimental conditions used in different interleaflet coupling studies likely give rise to apparently contradictory differences in phenomena that have been reported so far. While studies on SLBs have shown the *L*_*o*_ domain dominating the phase state of asymmetric membranes (26,32,33), asymmetric unsupported membranes, such as vesicles, show more variability where the *L*_*d*_ phase more commonly dominates but both *L*_*d*_ and *L*_*o*_ phase monolayers can dictate the bilayer properties. This is suggestive that the rigidity of solid supports favors the more rigid *L*_*o*_ phase dominating the properties of asymmetric membranes. This is consistent with previous reports of more ordered phases being favored in symmetric bilayer membranes within regions of local adhesion, where the free energy for formation of ordered membrane domains is lowered by mechanical suppression of thermal membrane undulations (34). Furthermore, in our current work, the only observation of the *L*_*o*_ phase dominating behavior is seen where the *L*_*o*_ phase GUV is under tension such that it does not visibly deform from its sphericity during hemifusion, giving rise to an interface with greater mechanical rigidity. Unsupported membranes may be more physiologically relevant to the behaviour of biomembranes within a cellular context, however mechanoresponsive mechanisms are known to play important roles in biological signalling processes. The ability of asymmetric membranes to change phase state in response to local changes in mechanical cues in their environment, such as during cell adhesion, the invasion of pathogens, or cytoskeletal interactions with the membrane could play a fundamental role in rapid cell signalling responses.

## CONCLUSION

Here we have demonstrated that 30 nm SiO_2_ NPs mediate fusion between *L*_*d*_ DOPC GUVs and *L*_*o*_ DPPC:Chol (7:3) GUVs and generate phase separated GUVs. Furthermore, we have shown that imaging of *L*_*d*_ GUVs and *L*_*o*_ GUVs undergoing hemifusion, allows real time visualization of asymmetric domain formation and interleaflet coupling. While membrane mechanical properties such as bending rigidity, lateral tension and curvature has been proposed to play a central part during membrane fusion (18,35-37), less attention has been paid to the role of the different lipid composition and biophysical properties of each membrane undergoing fusion. We find that transmembrane stresses generated in asymmetric membrane intermediates enhance the efficiency of full fusion between vesicles compared to symmetric membrane fusion in our previous studies (18). Additionally, we find that the *L*_*d*_ phase dominates the properties of bilayers with phase asymmetry; by comparison with broader findings in the literature, this may imply that the mechanical properties of the local environment could be an important parameter in determining the phase that dominates the local bilayer properties. Further investigations following this experimental approach using a wider range of vesicle compositions would provide additional valuable insight into the role of each leaflet’s composition, phase state and mechanical properties on interleaflet coupling in the mechanisms governing the lipid lateral organization and phase states within biological membranes.

## AUTOR CONTRIBUTIONS

MAP performed the experiments and the data analysis. MAP and PAB contributed to designing the experiments, discussion and writing the manuscript. PAB supervised the research.

## DECLARATION OF INTERESTS

The authors declare no competing financial interest.

## ACKNOWLEDGEMENTS

M. A. P. acknowledges the University of Leeds for funding through the Lowson Scholarship. The Zeiss LSM 880 with Airyscan inverted confocal micrsoscope was funded by the Wellcome Trust (WT104818MA).

## Notes

### Competing Interest Statement

The authors have declared no competing interest.

