## Supplemental Information for "Dynamics of asymmetric membranes and interleaflet coupling as intermediates in membrane fusion"

### Supplementary figures:

#### Figure S1.

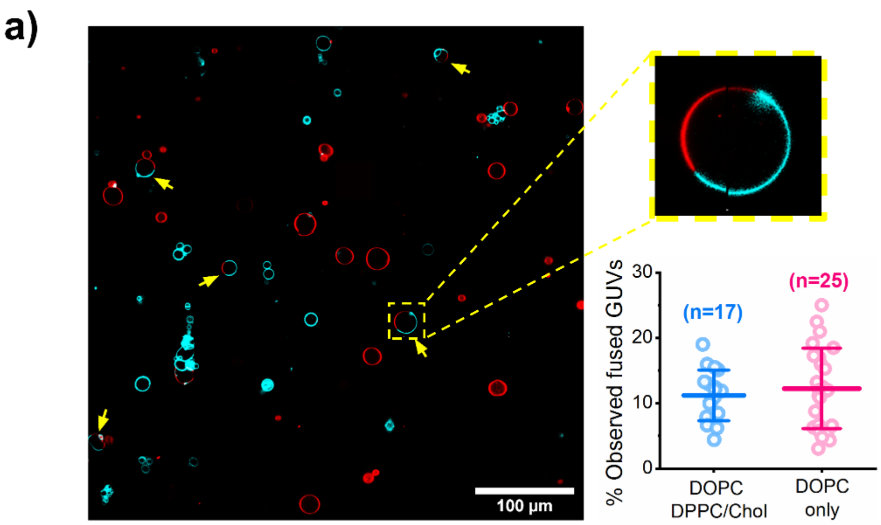


b)

| GUVs composition | Total fusion events recorded | Direct full fusion | Hemifusion-fusion | Gentle merging |
|---|---|---|---|---|
| DOPC only | 20 | 35% (n=7) | 30% (n=6) | 35% (n=7) |
| DOPC + DPPC:Chol (7:3) | 32 | 53% (n=17) | 47% (n=15) | 0% (n=0) |

**a)** Confocal microscopy tile scan after incubating DOPC GUVs and DPPC:Cholesterol (7:3) GUVs in a 1:1 volume ratio with 25 µg/ml $SiO_2$ NPs. Yellow arrows highlight the phase separated GUVs. Top left image shows a zoomed in phase separated GUV. Plot shows the distribution of the percentage of fused GUVs observed from tile scans in samples of DOPC GUVs mixed with DPPC:Chol (7:3) GUVs in a 1:1 volume ratio (blue) and in control samples of DOPC GUVs (magenta) where half of the GUVs are labelled with Rh-DOPE and the other half with DiO. Data are presented as mean ± SD, circles indicate the percentage of fused GUVs observed in each tile scan (number of tile scan images analysed indicated in the plot). In samples of DOPC and DPPC:Chol GUVs the proportion of observed fused (phase separated) GUVs was 11.25 ± 3.89 % and in samples of only DOPC GUVs the proportion of observed fused GUVs was 12.45 ± 5.93 %. The proportion of observed fused GUVs in the two conditions does not show statistical difference (one-way ANOVA with a $p < 0.5$ post-hoc Bonferroni test). Over 1000 individual GUVs were counted for each condition. **b)** Table shows the percentage of recorded fusion events corresponding to each fusion pathway described in our previous study (direct full fusion, hemifusion-fusion and gentle membrane merging) (1) when both GUVs fusion are composed of only DOPC and when one is DOPC and the other DPPC:Chol (7:3). While the percentage of the 3 pathways is very similar in DOPC-DOPC GUV fusion, the gentle membrane merging pathway is suppressed when one of the vesicles undergoing fusion is a liquid ordered DPPC:Chol GUV.

**Figure S2.**

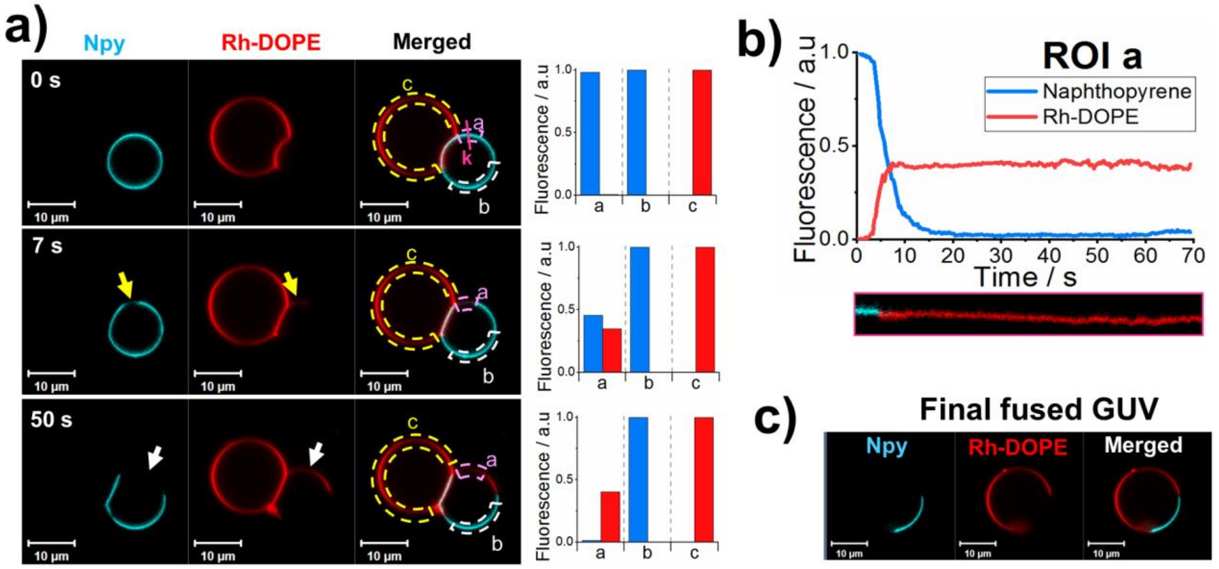


Asymmetric membrane domains and interleaflet coupling in hemifused GUVs. a) Confocal microscopy time-lapse images of GUVs during hemifusion. Naphthopyrene (Npy) (cyan) and Rh-DOPE (red) are used as $L_o$ and $L_d$ markers, respectively. Yellow arrows indicate regions of the membrane where both dyes are colocalised (asymmetric domains) and white arrows indicate regions of the membrane where the $L_o$ marker has been excluded from the membrane due to interleaflet coupling. Bar plots show the normalised fluorescence intensity measured in the ROIs "a","b" and "c" shown in the merged channel. Fluorescence intensity in "ROI b" and "ROI c" corresponds to the maximum intensity of Npy and Rh-DOPE, respectively, and the background is used as minimum intensity. "ROI a" shows an initial rise in Rh-DOPE intensity simultaneous to a reduction of Npy fluorescence to about half compared to its maximum (7 s micrograph). Micrographs at 50 s show a complete exclusion of Npy from that region of the membrane while the intensity of Rh-DOPE in "ROI a" remains close to 0.5. b) Normalised fluorescence intensity of Rh-DOPE (red line) and Npy (blue line) in "ROI a" against time. Kymograph shows fluorescence signal of the dyes over time at the pink line indicated as "k" in the 0 s micrographs in a). c) Phase separated GUV after full fusion.

**Figure S3.**

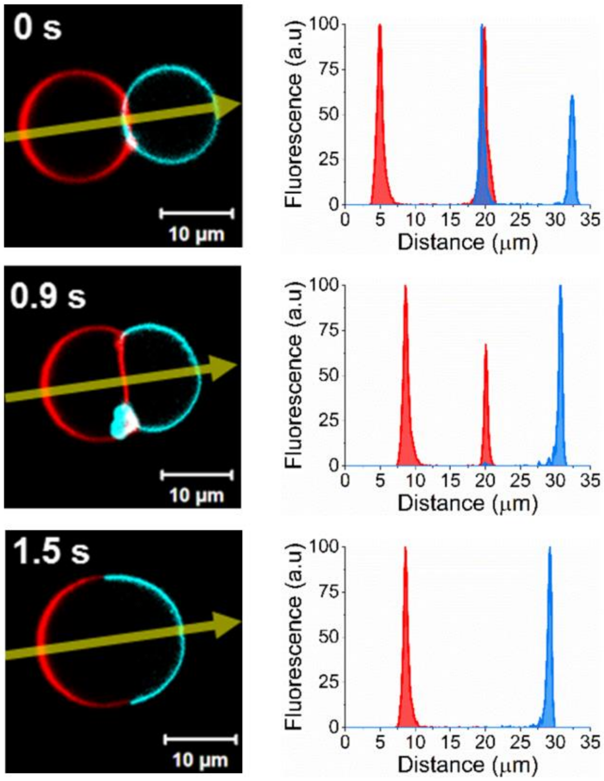


Confocal microscopy micrographs showing Naphthopyrene (Npy) exclusion from the contact region before fusion. Plots show the fluorescence intensity profile of Npy and Rh-DOPE across the yellow arrow indicated in the micrographs.

### Supplementary movies

**<u>File Name:</u> Supplementary Movie 1**
Description: Direct full fusion between a DOPC GUV labelled with Rh-DOPE (red) and a DPPC:Chol (7:3) labelled with Napthopyrene (cyan) mediated by 30 nm $SiO_2$ NPs. The fusion generates a phase separated GUV with a DOPC-rich *Ld* domain and a *Lo* domain composed mainly by DPPC and cholesterol.

**<u>File Name:</u> Supplementary Movie 2**
Description: Fusion between a DOPC GUV labelled with Rh-DOPE (red) and a DPPC:Chol (7:3) labelled with Napthopyrene (cyan) mediated by 30 nm $SiO_2$ NPs preceded by a hemifusion intermediate state. During hemifusion Rh-DOPE fluorescence arises in the *Lo* DPPC:Chol (7:3) GUV leading to a short-lived colocalisation of the dyes (asymmetric domain) followed by the exclusion of Napthopyrene from a region of the *Lo* GUV (interleaflet coupling).

**<u>File Name:</u> Supplementary Movie 3**
Description: Naphthopyrene exclusion from the contact region between a DOPC GUV labelled with Rh-DOPE (red) and a DPPC:Chol (7:3) labelled with Napthopyrene (cyan) before they fully fuse. Fusion is mediated by 30 nm $SiO_2$ NPs.

**<u>File Name:</u> Supplementary Movie 4**
Description: Rh-DOPE exclusion from the contact region between a DOPC GUV labelled with Rh-DOPE (red) and a DPPC:Chol (7:3) labelled with Napthopyrene (cyan). Full fusion is not observed.
